# An evidence-based approach to globally assess the covariate-dependent effect of MTHFR SNP rs1801133 on plasma homocysteine: a systematic review and meta-analysis

**DOI:** 10.1101/204487

**Authors:** Huifeng Jin, Haojie Cheng, Wei Chen, Xiaoming Sheng, Mark Brown, Junqiang Tian

**Author notes:** The authors contributed equally to the research (Huifeng Jin, Haojie Cheng). **Authors’ last names**: Jin, Cheng, Wei, Sheng, Brown, Tian. **Disclaimer**: none. **Clinical Trial Registry number and website**: none. **Abbreviations** FA: folic acid MTHFR: methylene tetrahydrofolate reductase SNP: single nucleotide polymorphism tHcy: total plasma homocysteine.

## Abstract

**Background:** The single nucleotide polymorphism (SNP) of the gene Methylenetetrahydrofolate Reductase (MTHFR) C677T (or rs1801133) is the most established genetic factor that increases plasma total homocysteine (tHcy) and consequently results in hyperhomocysteinemia. Yet given the limited penetrance of this genetic variant, it is necessary to individually predict the risk of hyperhomocysteinemia for a rs1801133 carrier.

**Objective:** We hypothesized that variability of this genetic risk is largely due to the presence of factors (covariates) that serve as effect modifiers and/or confounders, such as folic acid (FA) intake, and aimed to assess this risk in the complex context of these covariates.

**Design:** We systematically extracted from published studies the data of tHcy, rs1801133, and any previously reported rs1801133 covariates. The resulting meta-dataset was first used to analyze the covariates’ modifying effect by meta regression and other statistical means. Subsequently, we stratified tHcy data by the rs1801133 genotypes and analyzed under each genotype the variability of the risk resulted from the covariates’ confounding.

**Results:** The dataset contains data of 36 rs1801133 covariates that were collected from 114,448 subjects and 249 qualified studies, among which 6 covariates (sex, age, race, FA intake, smoking, and alcohol consumption) are the most frequently informed and therefore included for statistical analysis. The effect of rs1801133 on tHcy exhibits significant variability that can be attributed to effect modification and, to a larger degree, confounding by these covariates. Via statistical modeling, we predicted the covariate-dependent risk of tHcy elevation and hyperhomocysteinemia in a systematic manner.

**Conclusions:** we demonstrated an evidence-based approach that globally assesses the covariate-dependent effect of rs1801133 on tHcy. The results should assist clinicians in interpreting the rs1801133 data from genetic testing for their patients. Such information is also important for the public that increasingly receives genetic data from commercial services without interpretation of its clinical relevance.

## Introduction

MTHFR is a key enzyme in the one carbon unit metabolism that donates the methyl group through the methionine/homocysteine cycle. SNP rs1801133, a missense mutation from cytosine (C) to thymine (T) in exon 4 of MTHFR, results in a change in the amino acid sequence (alanine to valine) and consequently attenuates the catalytic activity (1). Since the first report in 1995 (2), it has been widely accepted that rs1801133 raises blood tHcy (free and protein bound homocysteine). Although definitive evidence is lacking regarding the causative role of elevated tHcy in various diseases, it has been generally considered as a suboptimal health condition. It is common practice for physicians to recommend tHcy-lowering measures, such as vitamin Bs, for rs1801133 carriers. As genotyping becomes commercially available, this practice is followed by public consumers who normally receive genetic data without professional interpretation.

However, this practice fails to take into consideration the individual variation of this genetic effect. In fact, the penetrance of this SNP is relatively small. For instance, the increase of the prevalence of hyperhomocysteinemia among TT and CT carriers in US population is only 6.5% and 1.8%, respectively, up from that of CC (14.4%) (3). From this perspective, a fundamental question remains unanswered: what are the measures for personalized evaluation of the risk of hyperhomocysteinemia for rs1801133 carriers?

The low penetrance of hyprserhomocysteinemia may be due to the influence of other factors (covariates) that are also known to affect plasma tHcy, including demographics (such as age, sex, race), nutrition (dietary intake and supplementation of vitamin B, choline, and protein), genetics (SNPs other than rs1801133), lifestyle (alcohol, smoking, and coffee), and medical treatments (4–6). These covariates may mask rs18011133’s effect as independent variables (confounders). More importantly, the covariates may interact with rs1801133 as effect mortifiers (or moderators) to change its effect on tHcy, thereby defying a uniform characterization of rs1801133 effect. This modifying effect has been demonstrated by several studies in which the effect of rs1801133 differs significantly as the status of the covariates change (3, 7–9).

Given the presence of multiple effect modifiers, we would expect to see a large variability of the tHcy-elevating effect of rs1801133 when it is viewed in the context of these modifiers. Nonetheless, since rs1801133 is only one of the factors that impact tHcy, the actual effect of rs1801133 on tHcy for any individuals who carry this variant is further complicated by the other covariates. Therefore, the risk of rs1801133-derived hyperhomocysteinemia has a two-fold complexity: effect modification and confounding. In this systematic review, we first made an effort to delineate the variability of this genetic effect in the context of its modifiers. Subsequently, we controlled for the genetic effect by genotype-stratifying the data before assessing the additional variability imposed by other tHcy determinants. With this sequential analysis, we attempted to address two closely related questions: 1) under what covariate-defined conditions is an individual carrying rs1801133 at risk for tHcy elevation? and 2) for the conditions under which a rs1801133-carrier is indeed at-risk for tHcy-elevation, what is his/her risk of hyperhomocysteinemia?

Notably, both of the above variabilities are dependent on not only the multitude of the covariates, but their combinations. As such, if a certain study determines the rs1801133 effect for a group of subjects with shared status on all covariates, it provides information to only one condition of the total combinations; and therefore, it would be ideal yet conceivably unattainable for a single research project sufficiently powered to map out the risks under all conditions. Nonetheless, global assessment can be achieved when methodologically compatible studies that investigated different conditions are aggregated following the stringent procedure of systematic review. In light of this, we catalogued available metadata from publications that include rs1801133 genotypes, tHcy, and the status of the relevant covariates. Such a meta-dataset allowed us to systematically characterize the covariates’ influence on rs1801133-derived tHcy-elevation and homocysteinemia.

## Methods

### 1. Data sources and search strategy

We first conducted a systematic literature search to gather data from available publications that reported the effect of rs1801133 on tHcy. Related publications by the date of December 17th, 2016 were identified by searching the databases (PubMed and Scopus) using 3 groups of search terms: 1) the gene (MTHFR), 2) the SNP (rs1801133), and 3) the trait (tHcy). Detailed search terms (including MESH terms in PUBMED) are provided in the Supplemental Material #1. Manual searches of references supplemented the electronic search. The search and review were conducted by two independent researchers (HJ and HC), and any disagreement was resolved by discussion with a third reviewer (JT). This study was registered at Research Registry (http://www.researchregistry.com/browse-the-registry.html#registryofsystematicreviewsmeta-analyses/) with the registration number: reviewregistry328.

### 2. Study eligibility and selection

As shown in Figure 1, All of the above search results (n = 3,671) were assessed according to the inclusion and exclusion criteria outlined in Supplemental Material #2. In brief, we included any clinical (interventional or Observational) studies that investigated the general population and provided the subjects’ data of tHcy and rs1801133. The studies were first screened by title, abstracts, and database annotations to exclude duplicates and non-related articles, such as non-human study, case report, review article, letter to the editor. We also excluded non-English articles and conference publications (posters and abstracts). Subsequently, we reviewed the full texts to exclude the studies in which: 1) the data had been previously reported in another publication; 2) the subjects were recruited exclusively from a patient population, such as the case group in a case-control study; 3) the genotype/tHcy data were incomplete or missing; 4) the tHcy data were provided as median instead of mean; 5) tHcy data were adjusted by variables include in this dataset; 6) studies that recruited family members. A complete list of included articles (n = 249) are shown in Supplemental Material #3.

**Figure 1.**
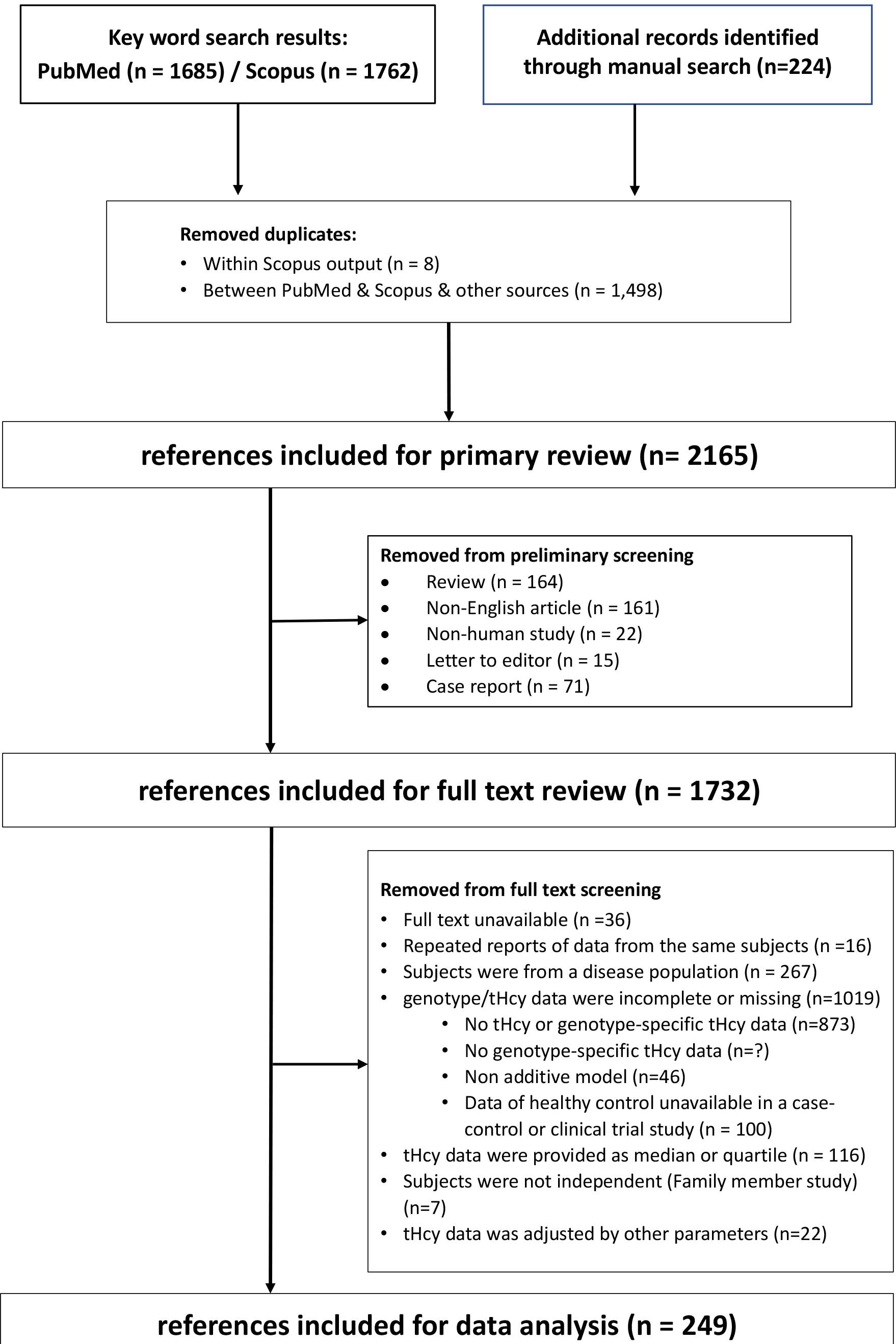
Overflow of Online Search Results

### 3. Covariate categorization and data extraction

Based on our knowledge, apart from rs1801133, 36 other tHcy determinants had been reported in literature that include age, gender, ethnicity, serum FA concentration, smoking, and alcohol drinking, other tHcy-relevant SNPs, physical activity, and the intakes of tHcy-relevant vitamins (folate, vitamin B2, B6, and B12), choline, and protein. Among these variables, FA serum concentration was excluded as an independent variable for covariate analysis due to its collinearity with FA intake (data not shown). To standardize data extraction for these variables, we categorized each variable as detailed in Supplemental Material #4.

Since each of the included studies may contain one or more experimental group (named “Observation”) that is differentiated by the covariate status, we collected a total of 353 Observations from the 249 articles, and extracted data for each of the Observations that include sample size, sampling date and country, genotypes, tHcy, and the status of 35 associated variables. If for a certain variable the subjects in an experimental group have mixed status (such as containing both males and females for sex), it is denoted as “mixed” for this Observation. If information is not provided in a study for a certain covariate, it is denoted as “unknown”.

### 4. Risk-of-bias Analysis

We examined the risk-of-bias with each Observation using Item Bank on Risk of Bias and Precision of Observational Studies (10) and Cochrane Handbook for Systematic Reviews of Interventions tool (11). Since our analysis was based on the Observations instead of studies, we adapted the above assessment tools to reflect the potential within-Observation bias among the 3 rs1801133 genotypes in selection, detection, performance, and reporting. We also considered the selection bias resulted from the deviation of the subjects from the general population, such as sampling from hospital patients and the healthy worker’s bias. The scoring method for these biases are detailed in Supplemental Material #5. Clinical trials for which the tHcy data were obtained at baseline are considered as cross-sectional studies.

### 5. Data synthesis and statistical analysis

Given the skewed distribution of tHcy in many of the reports, tHcy data were log-transformed for analysis. All analyses were conducted using Comprehensive Meta-Analysis (CMA v3.0, Biostat Inc., Englewood, NJ) or SAS (v9.4, SAS Institute Inc., Cary, NC).

Where applied, the “mixed” and “unknown” data were regarded as independent categories within a variable. On rare occasions, “unknown” was combined with another category of one covariate if the post hoc analysis indicated non-significant difference between these two. To ensure the quality of the analysis, if the total Observation number was < 6 and subject number < 650 for one category, it was excluded from analysis. We also excluded the variables that have unknown status for more than 270 Observations or less than 25,000 subjects. This resulted in the retention of six variables: sex, age, race, folate intake, smoking, and drinking.

For meta-analysis of the overall rs1801133 effect on tHcy, a pooled weighted mean difference (WMD) with 95% confidence interval (CI) was calculated using random-effect model. We assessed the publication bias of this effect using Begg’s funnel plot and Egger’s test (12), and the heterogeneity between the Observations using the I^2^ and chi-square based Q statistic (13). The covariates’ modifying and confounding effect were analyzed using regression models with ΔHcy (CT vs. CC and TT vs. CC) and tHcy as the outcome, respectively.

We also used hierarchical matrix (14) to analyze how the covariates modified the rs1801133 effect on tHcy. In brief, we sequentially stratified the data of rs1801133 effect (ΔtHcy of TT vs. CC and CT vs. CC) by each of the variables, which resulted in an array of cells each containing data collected under a specific multi-variable-defined condition. We then followed the random effect model to aggregate the data in cells where more than one Observation was entered. Power analysis was conducted to assess the number of subjects required for a power of >80%, alpha level of 5%, and minimal effect size as the WMDs obtained from the dataset. To ensure this power level, cells with a sample size of less than 115 (TT vs. CC) and 860 (CT vs. CC) were excluded from further analysis.

## Results

### 1. The rs1801133 effect on tHcy in the general population is highly heterogeneous

The systematic review of studies on rs1801133 and tHcy resulted in a dataset of 249 publications that contain relevant information from 353 experimental groups (hence 353 Observations). The total subjects (n=114,448) consisted of CC (n=50,296, 43.95%), CT (n=49,024, 42.84%), and TT (n=15,128, 13.22%) carriers, and the reported tHcy level for the 3 genotypes ranged from 5.1 to 38.6 uM (CC), 5.5 to 29.7 uM (CT), and 3.15 to 92.4 uM (TT), respectively. The Observations were then classified based on the categories of each of the 34 variables in our dataset (Figure 2A and Supplemental Material #6).

**Figure 2.**
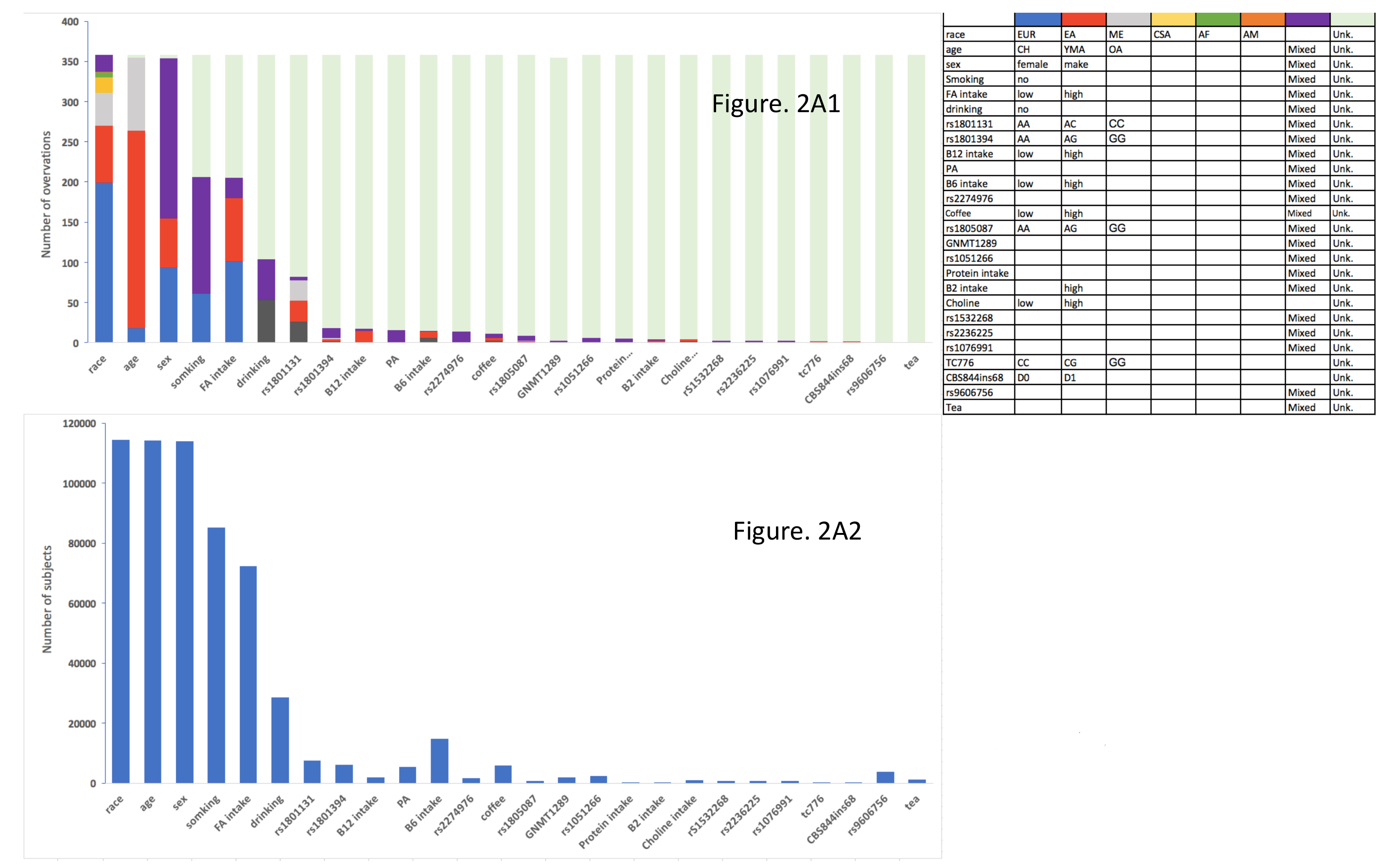
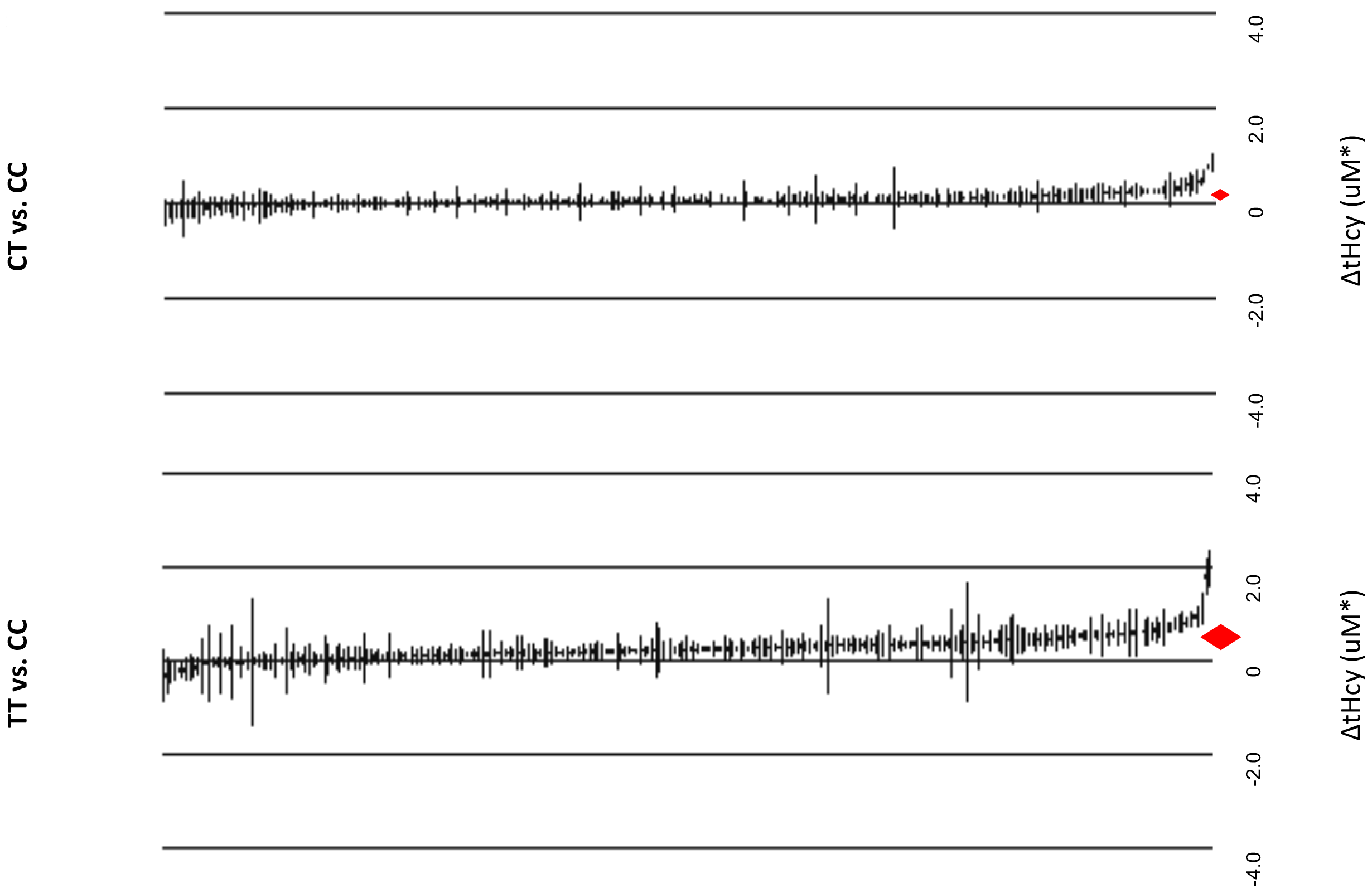
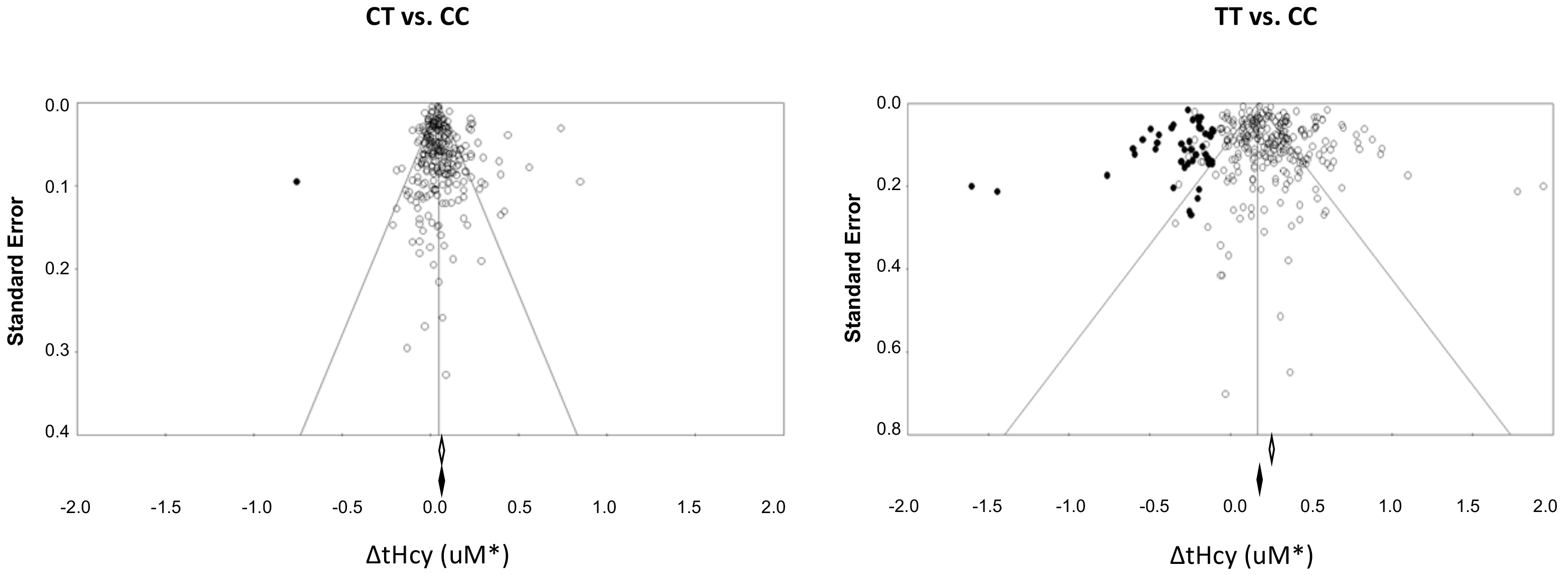
Overview of the meta-dataset. A, Number of Observations (A1) and subjects (A2) for tHcy-relevant variables. The variables without any Observation (n=7) was not shown. B, forest plot of the tHcy-elevating effect of rs1801133 (CT and TT, respectively). Heterogeneity for CT vs. CC: Tau^2^ = 0.0, Q = 1707, df = 277, I^2^ = 84%, p < 0.0001; and TT vs. CC: Tau^2^ = 0.0, Q = 3760, df = 272, I^2^ = 0.93, p < 0.0001. C, the Egger plot for CT and TT effect on tHcy. Egger’s Regression Intercept for CT: t-value=2.2, df=276, p=0.03; and TT: t-value=2.4, df=271, p=0.02. Solid dots indicate the hypothetical additional data required to avoid the effect bias. EUR, European; EA, East Asian; ME, Middle East; CSA, Central Southeast Asian; AF, African; AM, Amerindian; CH, child; YMA, young and middle-aged adult; OA, Old Adult; PA, physical activity. B2, B6, and B12 for vitamin B2, B6, and B12. *, data are on log(e) scale.

Since the status of some variables were often not disclosed, there was a large presence of the “unknown” category in these variables (Figure 2A1). Accordingly, the subject numbers that contribute to analyzing these variables’ effects were limited (Figure 2A2). In this respect, sex, race, and age, folate intake, smoking, and drinking were the most informed variables and were the variables included in subsequent statistical analyses. In addition, the category-wise distribution of data within each variable was skewed by available studies. For instance, compared with other race categories, the data of race is unavailable for “Oceanian” and limited for Amerindian (n=1 Observation) and African (n=6 Observations).

Next, we examined any potential sources of bias in each Observation that may skew the tHcy level or the rs1801133-tHcy association (Supplemental Material #7). In about 1/3 of the Observations, the sampling was conducted via convenient methods, such as that from hospital workers or outpatients, which may cause deviation of the subjects from the intended general population. The lack of covariate specification is a prominent risk for selection bias among the 3 genotypes. In addition, over half of the clinical trials (mainly FA supplementation) did not provide information for proper randomization and blinding of the investigators and the subjects. With this dataset, we first determined the overall tHcy-elevating effect of rs1801133 (as defined by the log_e_ mean difference of tHcy of the CT and TT genotype carriers over those of CC). This effect is highly significant (P < 0.0001) for both CT (0.062 uM, 95%CI = 0.051 – 0.072 uM) and TT (0.250 uM, 95% CI = 0.226 – 0.274 uM) genotype. Figure 2B presents the forest plots of the results with weighted overall means. Notice that despite the high significance level, the reported effects vary significantly as demonstrated by the marked homogeneity index (I^2^ = 84% for CT and 93% for TT, for both P < 0.0001), suggesting the presence of effect modifiers. The funnel plots of the overall results indicate a bias towards a tHcy-increasing effect for both TT and CT genotypes (Figure 2C), suggesting the presence of publication bias.

### 2. The rs1801133 effect on tHcy is modified by multiple covariates

Given the large heterogeneity shown in Figure 2B, we determined if this tHcy-elevating effect is modified by covariates. Meta regression analysis indicate that sex, race, and FA intake are strong effect modifiers (P < 0.0001) for both CT and TT genotypes (Figure 3A). Specifically, the tHcy-elevating effect is decreased in females (vs. males) and in people consuming FA supplement or FA-fortified food. Individuals of Central and South Asian and Middle East tend to enhance the effect than other races. Detailed information of the effect sizes of each variable is available in Supplemental Material #8. Next, we assessed how these modifiers collectively alter the effect of rs1801133. Using the modeling results (the coefficients for the variables and intercept), we predicted this effect under each variable-defined condition (a specific combination of the variables). When viewed across the variable-defined spectrum, the rs1801133 effect is manifest in most, but not all, conditions (84.1% for CT and 94.7% for TT, respectively; Figure 3B and Supplemental Material #9), indicating the variability of this effect. In addition, the effect size varies considerably from −0.08 to 0.19 uM (CT) and −0.07 to 0.52 uM (TT) on log_e_ scale. This variation may therefore explain in part the heterogeneity observed in Figure 2B.

**Figure 3.**
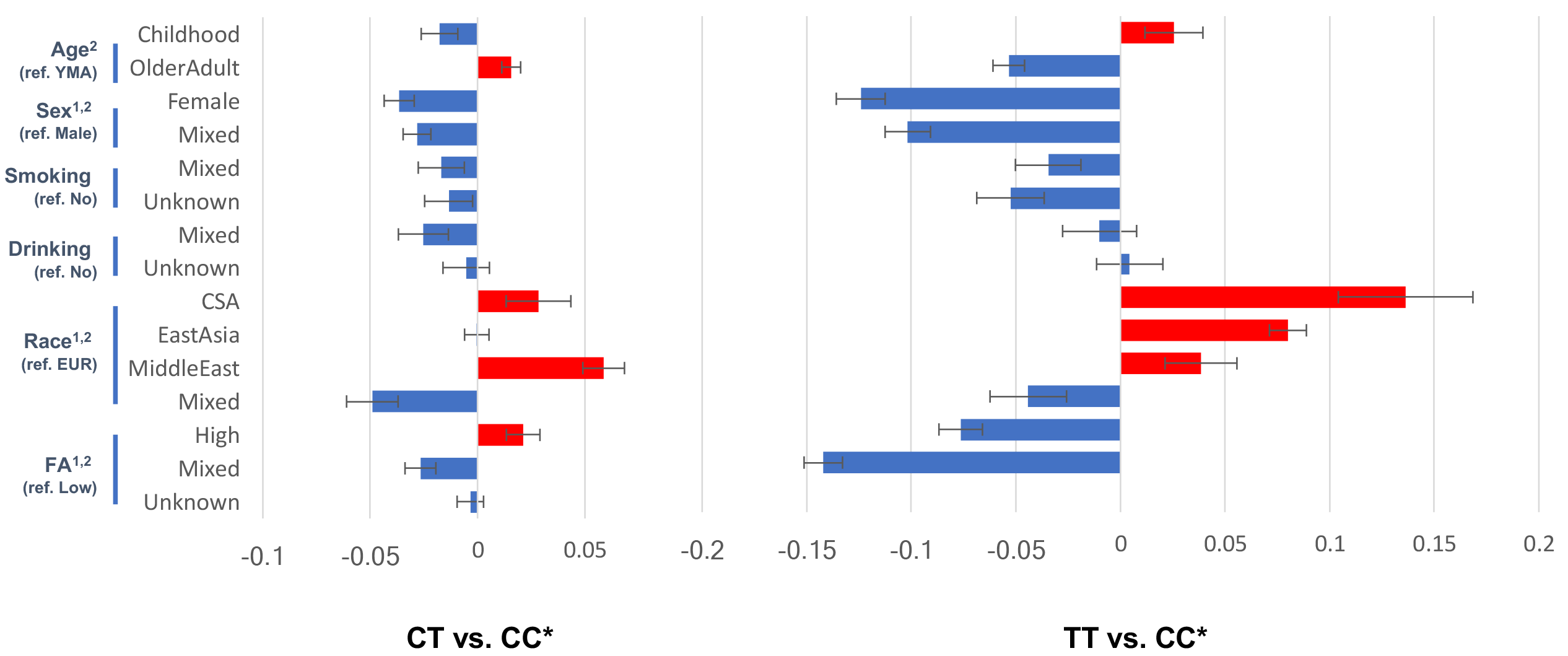
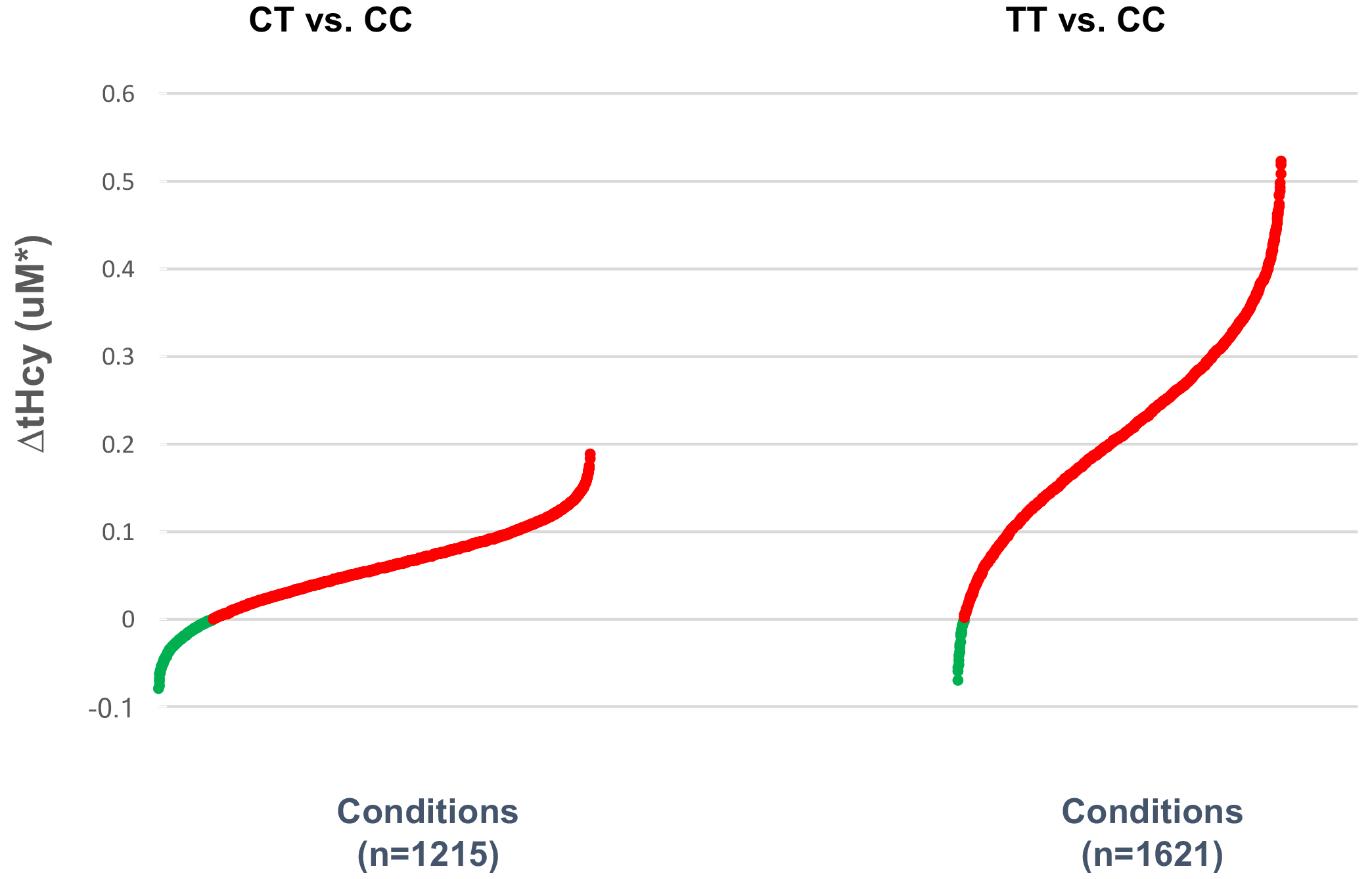
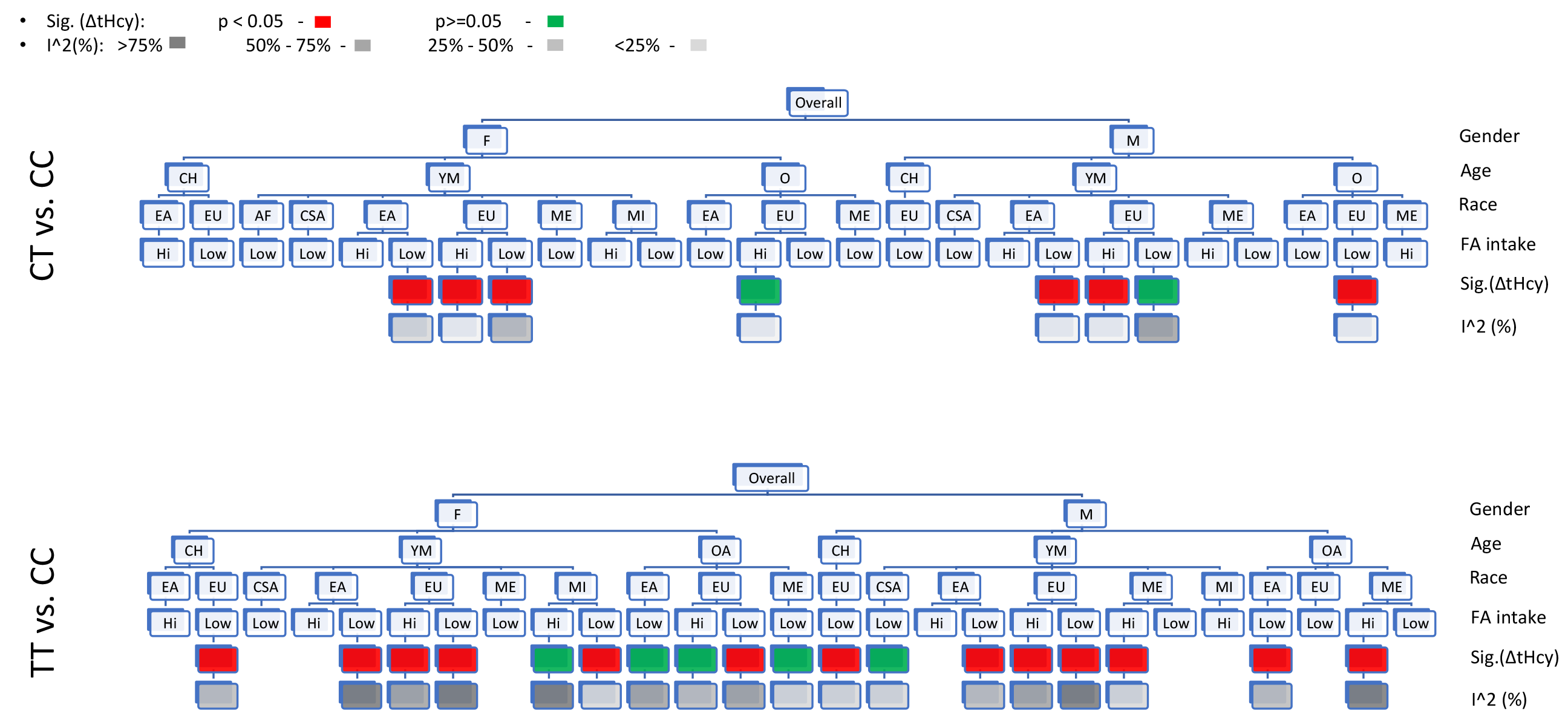
Statistical analyses of the modifiers of the rs1801133 effect on tHcy. A, Meta regression analysis. *, data are on log(e) scale. A1, diagrammatic presentation of the variables’ modifying effect. ^1^ and ^2^, p < 0.0001 for variable’s overall effect for CT vs. CC and TT vs. CC, respectively. Bars represent standard error. ref, reference category; For TT vs. CC, the FA category “unknown” was combined with “low” after post hoc test. A2, tHcy-elevating effect of rs1801133 (TT and CT) across variables’ combinatorial conditions. Green and red dots represent conditions with predicted ΔtHcy < 0 and >0, respectively. B, Hierarchical matrix analysis. The diagrams illustrate the sequential data stratification. Cells not populated with data are not shown, and the cells without sufficient power are not analyzed for significance and I^2^. EUR, European; EA, East Asian; ME, Middle East; CSA, Central Southeast Asian; M, mixed racial background; CH, child; YMA, young and middle-aged adult; OA, Old Adult.

Since the Observations with “unknown” and “mixed” variables reduced the power of the regression analysis, we next used a more stringent method (hierarchical matrix) to analyze the modifiers’ effect. In this method, we first removed any Observations with “unknown” or “mixed” status from the dataset, and stratified the remaining data into an array of cells (n = 18 for TT and 8 for CT), each representing a specific combination of the variables (Figure 3C). As expected, the stratification decreased the heterogeneity index (I^2^) of tHcy in each cell as compared with overall data (Figure 2B). Statistical analysis indicates that, in keeping with the results from meta regression, the rs1801133 effects not only vary significantly with the covariates’ status (Supplemental Material #10), but also become insignificant for part of the cells (5 out of 18 for TT and 2 out of 8 for CT; Figure 3C). The results thus indicate that a non-discriminant risk prediction based solely on rs1801133 genotype is overly simplistic.

### 3. The rs1801133 effect on tHcy is confounded by multiple covariates

Next, we sought to assess the influence of rs1801133’s covariates that confounded the its effect on tHcy. For this purpose, we first controlled for the covariates’ modifying effect by stratifying the tHcy data with the three genotypes of rs1801133, and then analyzed under each genotype the status of tHcy in association to the variables. As reported, tHcy was significantly affected by age, sex, race, FA intake, smoking, and drinking across all three genotypes (P < 0.0001, Figure 4A, Supplemental Material #11), among which smoking (even as mixed status) and race are the most pronounced confounders. It is notable that the confounding effect of the covariates is in general much larger in effect size than that of modifiers (Figure 3A1). Nonetheless, none of the variables alone dictates the incidence of hyperhomocysteinemia.

**Figure 4.**
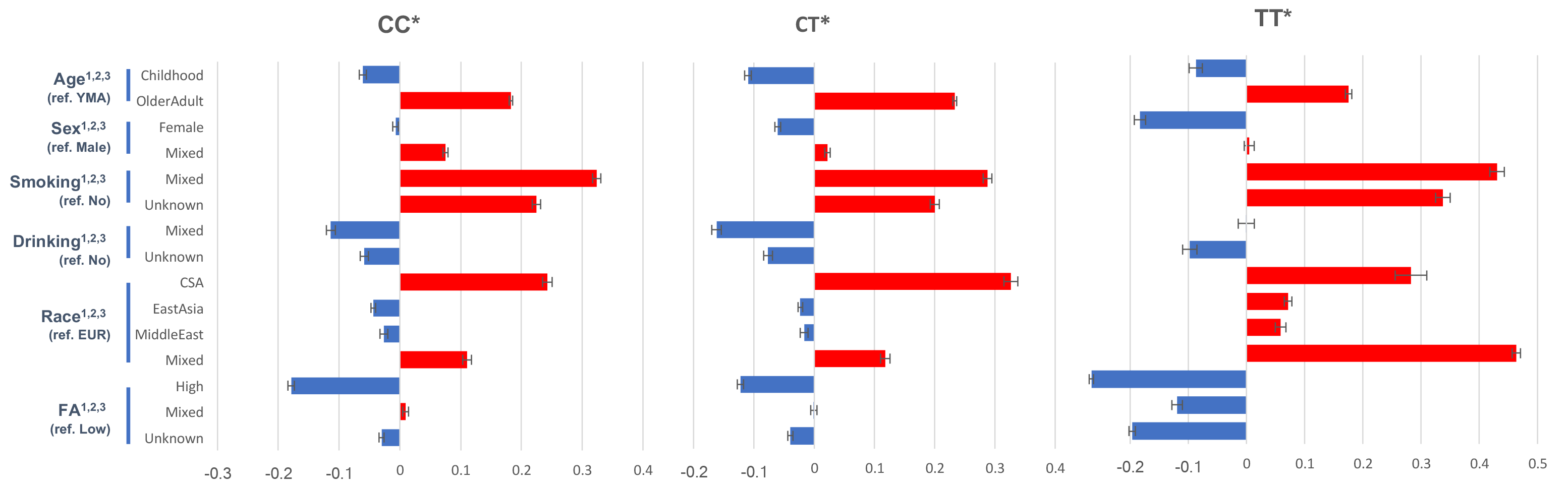
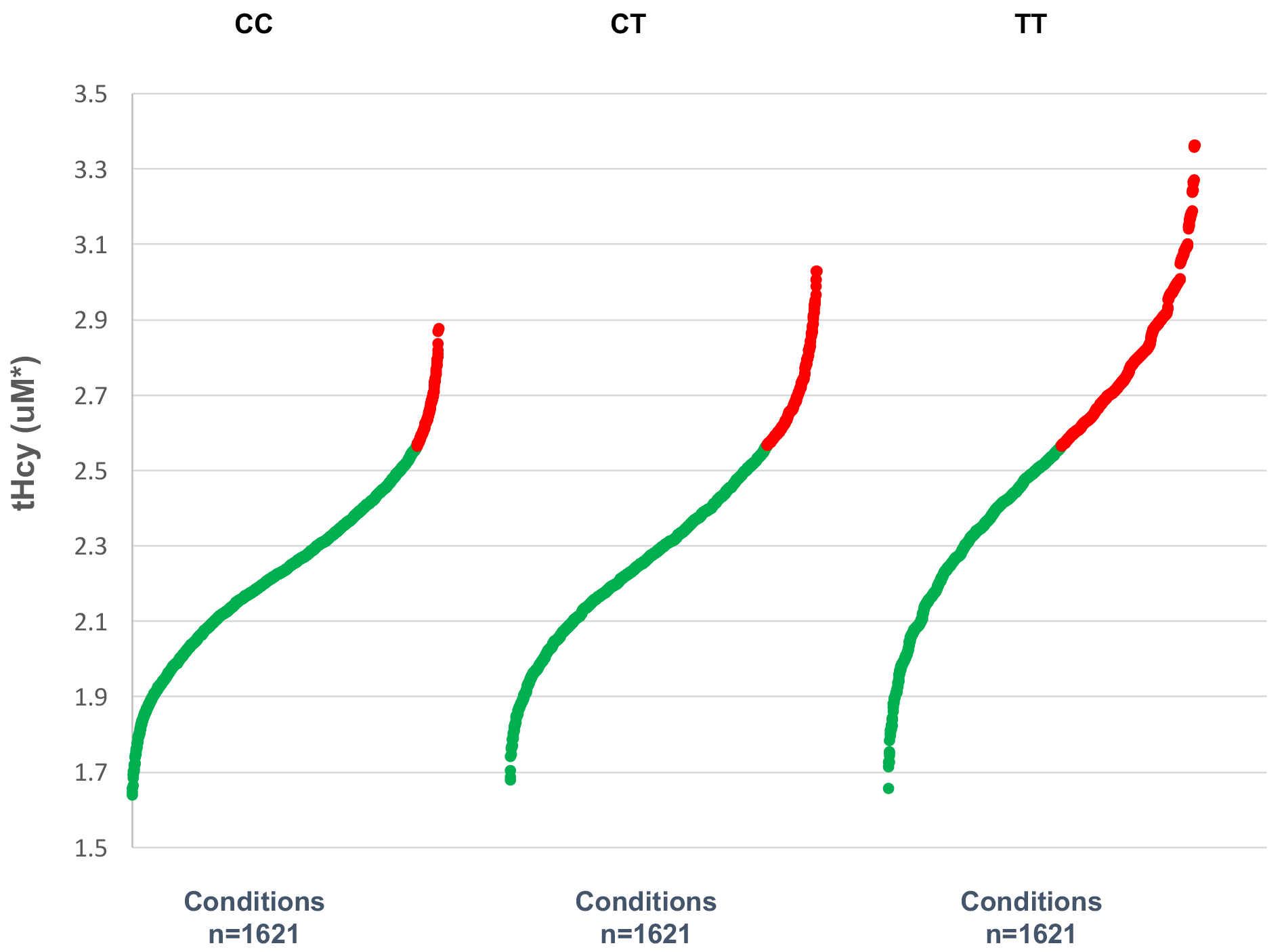
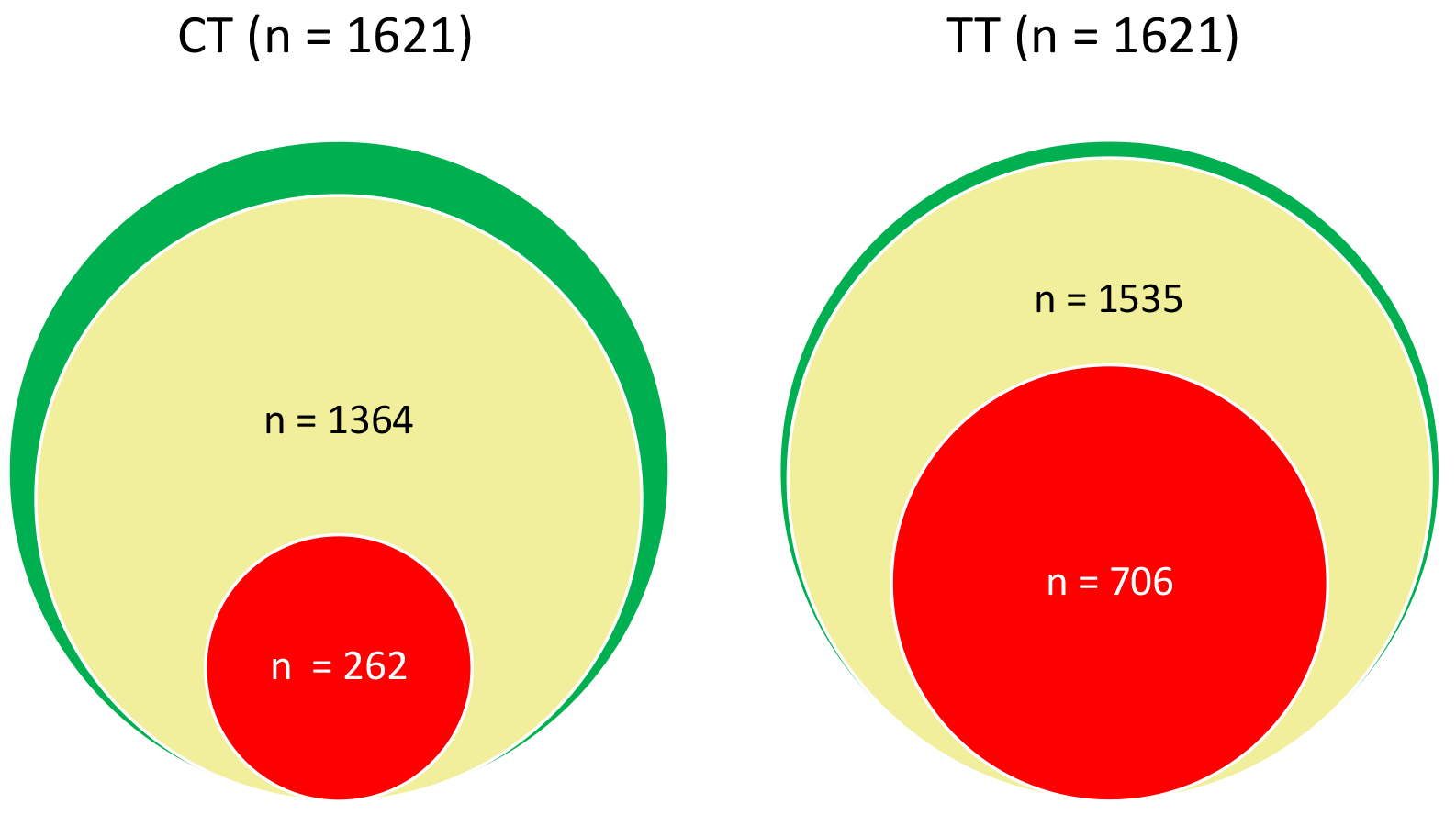
Regression analysis of the confounders of the rs1801133 effect on tHcy. A, diagrammatic presentation of the covariates’ confounding effect on tHcy. Bars represent standard error; ref, reference category; *, on log(e) scale; ^1^, ^2^, and ^3^, p < 0.001 for variable’s overall effect for CC, CT, and TT, respectively. EUR, European; CSA, Central Southeast Asian; YMA, young and middle-aged adult. B, Risk of hyperhomocysteinemia across different covariate-defined conditions for each rs1801133 genotype. Red and green dots represent conditions with Log transformed tHcy above (hyperhomocysteinemia) and below (normohomocysteinemia) Log_e_13 uM, respectively. C, overview of the covariant-dependent variability of rs1801133 effect on tHcy. The green spheres represent the total number of covariate-defined conditions, the yellow spheres represent the conditions with tHcy elevation, and the red spheres represent conditions that exhibit both tHcy elevation and hyperhomocysteinemia.

Next, Using the results obtained from the above model, we predicted the tHcy values under each covariate-defined condition. The values were then classified as “hyperhomocysteinemia” or “normohomocysteinemia” based on a conventional cut-off of 13 uM (Figure 4B and Supplemental Material #12). As expected, the number of hyperhomocysteinemic conditions increases from CC to CT and then to TT. Importantly, even under TT, the number of hyperhomocysteinemia is less than half of the total conditions, indicating again the importance of covariate-dependent risk prediction.

Finally, in Figure 4C we summarize the covariate-defined conditions that are at risk for tHcy elevation and subsequently hyperhomocysteinemia. First, among a total of 1,621 covariate-defined conditions each for CT and TT, 84.1% (CT) and 94.7% (TT) of which exhibit tHcy elevation. Next, within the conditions with tHcy elevation, 19.2 % (CT) and 46.0% (TT) of which exhibit hyperhomocysteinemic level of tHcy.

## Discussion

In this study, we utilized published data to assess the variability of the genetic variant (rs1801133)’s effect on tHcy in a covariate-dependent manner. We demonstrated systematically that this genetic risk, either in its form of tHcy elevation or hyperhomocysteinemia, is not only dependent on a number of covariates, but also on the combination of these covariates. For instance, for a female senior East Asian with low FA intake, TT genotype does not pose a risk of tHcy increase. However, the risk becomes significant if the sex status changes to a male or the race to European (Figure 3B). Such findings highlight the caution that needs to be exercised when using genetic data for predicting an individual’s tHcy risk.

There is a large body of publications regarding rs1801133 that has accumulated since the first report of this genetic risk in 1995 (2). In this study, we collected 249 relevant articles that amounts to 353 experimental groups (Observations) based on the status of 34 covariates. However, as shown in Figure 2A, none of these studies provided complete information for these covariates. Even for the six covariates that are most frequently informed and therefore included in our regression analyses (age, sex, race, supplement, smoking, and drinking), only about a quarter of the studies provided complete information. In addition, the covariates’ status in some study groups was indicated but indiscriminant (e.g., containing both sexes). As a result, there is a large presence of missing (“unknown”) and “mixed” data in our dataset, which increased the risk for selection bias and decreased the power of the statistical modeling. Another power issue comes from the age designation where the study group were assigned to an age category by the average age, leaving the possibility of mismatch for some individuals. Nevertheless, the modeling revealed strong modifying (Figure 3) and confounding (Figure 4) effects, which explains the high variability of reported data on rs1801133-driven tHcy elevation and hyperhomocysteinemia.

Given the 6 covariates (age, sex, race, supplement, smoking, and drinking), we were able to predict that the risk of hyperhomocysteinemia occurred in only part of the covariate-defined conditions. This prediction is based on statistic modeling, which allows us not only to integrate multiple studies conducted under one condition but also to interpolate from studied conditions to non-studied conditions. The predicted results are yet to be validated by future studies. Nonetheless, this evidence-based approach provides a reference for clinicians for whom currently there is no clinical guidance on reporting this risk in a personalized way. More relevantly, such information may help the general public that receive genotype data through commercial service but without professional guidance. Given that rs1801133 is a relatively common SNP, with an overall prevalence in our dataset of 43% for the heterozygous (CT) and 13% for homozygous (TT) carriers, a large number of people could potentially benefit from this approach of personalized risk prediction.

Since we stress that only part of rs1801133 carriers are at risk for tHcy elevation and hyperhomocysteinemia (Figure 3 and 4), we paid attention to the level of stringency against false negative finding (type II error) of the risk. Power analysis was conducted in the matrix analysis (Figure 3B) to ensure that the non-significant cells were not due to insufficient sample size. In addition, given the publication bias towards increasing overall tHcy-elevating effect of rs1801133 (Figure 2C), the estimated results from our analyses of this effect may be overestimated but lends more support to our conclusion.

Overall, our data indicate the importance of reporting the rs1801133 risk on tHcy in a covariate dependent way. From this perspective, we propose that future studies obtain subjects’ status of relevant covariates, and that attempts are made to utilize homogeneous subject populations with regards to these covariates. Such studies will not only reduce effect modification and confounding, but allow risk integration in later meta analyses with increased power. We appreciate Man li from the University of Utah in her inputs throughout this project, and Mark Levy from USANA Health Science in reviewing this article and providing valuable comments and suggestions.

This research is funded by USANA Health Science, Inc.

## Authors’ contributions

1. Junqiang Tian designed research (project conception, development of overall research plan, and study oversight); Xiaoming Sheng contributed to the study design and statistical analysis
2. Huifeng Jin, Haojie Cheng, and Wei chen conducted research (hands-on conduct of the experiments and data collection);
3. Huifeng Jin and Haojie Cheng analyzed data or performed statistical analysis;
4. Junqiang Tian wrote paper;
5. Junqiang Tian had primary responsibility for final content;

All authors read and approved the final manuscript.

